# GGCX promotes Eurasian avian-like H1N1 swine influenza virus adaption to interspecies receptor binding

**DOI:** 10.1101/2024.03.18.585644

**Authors:** Jiahui Zou, Meijun Jiang, Rong Xiao, Huimin Sun, Hailong Liu, Thomas Peacock, Shaoyu Tu, Tong Chen, Jinli Guo, Yaxin Zhao, Wendy Barclay, Shengsong Xie, Hongbo Zhou

## Abstract

The Eurasian avian-like (EA) H1N1 swine influenza virus (SIV) possesses the capacity to instigate the next influenza pandemic, owing to its heightened affinity for the human-type α-2,6 sialic acid (SA) receptor. Nevertheless, the molecular mechanisms underlying the switch in receptor binding preferences of EA H1N1 SIV remain elusive. In this study, we conducted a comprehensive genome-wide CRISPR/Cas9 knockout screen utilizing EA H1N1 SIV in porcine kidney cells. Knocking out the enzyme gamma glutamyl carboxylase (GGCX) reduced virus replication *in vitro* and *in vivo* by inhibiting the carboxylation modification of viral haemagglutinin (HA) and the adhesion of progeny viruses, ultimately impeding the replication of EA H1N1 SIV. Furthermore, GGCX was revealed to be the determinant of the D225E substitution of EA H1N1 SIV, and GGCX-medicated carboxylation modification of HA 225E contributed to the receptor binding adaption of EA H1N1 SIV to the α-2,6 SA receptor. Taken together, our CRISPR screen has elucidated a novel function of GGCX in the support of EA H1N1 SIV adaption for binding to α-2,6 SA receptor. Consequently, GGCX emerges as a prospective antiviral target against the infection and transmission of EA H1H1 SIV.

## Introduction

Influenza A virus (IAV) is a highly contagious respiratory pathogen responsible for annual epidemics and sporadic pandemics, causing significant morbidity worldwide ^1,2^. Due to the simultaneous expression of the avian-like α-2,3 and human-like α-2,6 sialic acid (SA) receptor, pigs can serve as intermediate hosts between birds and humans, facilitating the adaptation of avian influenza viruses (AIV) ^3,4^. The Eurasian avian-like (EA) clade 1C H1N1 swine influenza viruses (SIVs) originated from avian sources in 1979, subsequently spread through pig populations in Europe and Asia and causing sporadic human infections ^5,6^. Over prolonged evolution, the receptor binding preferences of EA H1N1 SIV have switched from the α-2,3 SA receptor to the dual binding of α-2,3 SA and α-2,6 SA receptor, and some recent isolates of EA H1N1 SIV have demonstrated greater propensity to bind to α-2,6 SA receptor ^7,8^. Moreover, after the global spread of the 2009 H1N1 pandemic and rapid reverse-zoonosis of this virus back in to pigs, reassortants between EA H1N1 SIV with the H1N1pdm09, arose widely, some of which exhibit effective transmission through respiratory droplets in ferrets ^9,10^, placing these viruses as potential instigators of the next influenza pandemic ^11,12^.

The emergence of an antigenically novel virus capable of efficiently infecting and transmitting between humans creates a scenario where EA H1N1 SIV could cause a pandemic ^13–15^. During the evolution of EA H1N1 SIV, adaptive mutations have arisen to overcome barriers between species, and specific viral substitutions have been identified as being critical for the transmission of the virus within mammalian populations. The E190D and D225E substitutions in HA are well characterized as altering sialoglycan specificity ^16–18^. Viruses with 225E in HA replicated faster than those with 225G due to differences in assembly and budding efficiency, possibly because the HA 225 mutation alters the salt bridge structure between amino acids D225 and K222, resulting in the receptor binding switch ^19,20^. The complex interplay between virus and host factors determines the adaptation of EA H1N1 SIV to different host species ^21,22^. However, the critical factor that contributes to HA 225E adaptation remains unknown. Therefore, identifying the host factors required for EA H1N1 SIV infection and exploring the potential host factors that drive adaptive substitutions contribute to elucidate the mechanisms underlying interspecies transmission and facilitate the development of targeted interventions aimed at disrupting its transmission pathways.

Using a CRISPR/Cas9-based high-throughput loss-of-function screening approach, we are able to provide a functional genomics resource critical for understanding the host factors involved in EA H1N1 SIV infection in pigs. Here, vitamin K-dependent gamma-carboxylase (GGCX) was identified for the first time as an essential host factor for EA H1N1 SIV infection. Our results show that GGCX catalyzes the carboxylation modification of viral HA protein, crucial for progeny virus binding to α-2,6 SA receptors. These findings suggest that GGCX is a major underlying driver of D225E substitution of EA H1N1 SIV and catalyzes the carboxylation modification of viral HA 225E, which determines the receptor binding preference of the EA H1N1 SIV, and may serve as a potential target to prevent EA H1H1 SIV replication and cross-species transmission.

## Results

### GGCX is required for efficient EA H1N1 SIV replication as identified by an unbiased genome-wide CRISPR knockout screen in pigs

To identify the host factors necessary for EA H1N1 SIV infection, we conducted a genome-wide CRISPR screen in Cas9-expressing porcine kidney-15 cells (PK-15-Cas9) following an established procedure (Fig. 1a) ^23^. Both parental cells and cells expressing version 1.0 of the PigGeCKO single guide RNA (sgRNA) libraries were infected with EA H1N1 SIV (A/Swine/HuBei/221/2016, HuB/H1N1) at a multiplicity of infection (MOI) of 0.01 and performed live/dead screens. Following four rounds of screening, we subjected the surviving cells from the second, third, and fourth rounds of challenge to high-throughput sequencing, then analyzed and ranked candidate genes using the model-based analysis of the genome-wide CRISPR/Cas9 knockout (MAGeCK) program ^24^. Enrichment of 14 genes (RBM6, SLC35A1, GGCX, ADD1, LOC100516036, GPCPD1, CMAS, ST3GAL4, etc) was observed in the second, third, and fourth rounds (Supplementary Table 1 and Extended Data Fig. 1).

**Figure 1.**
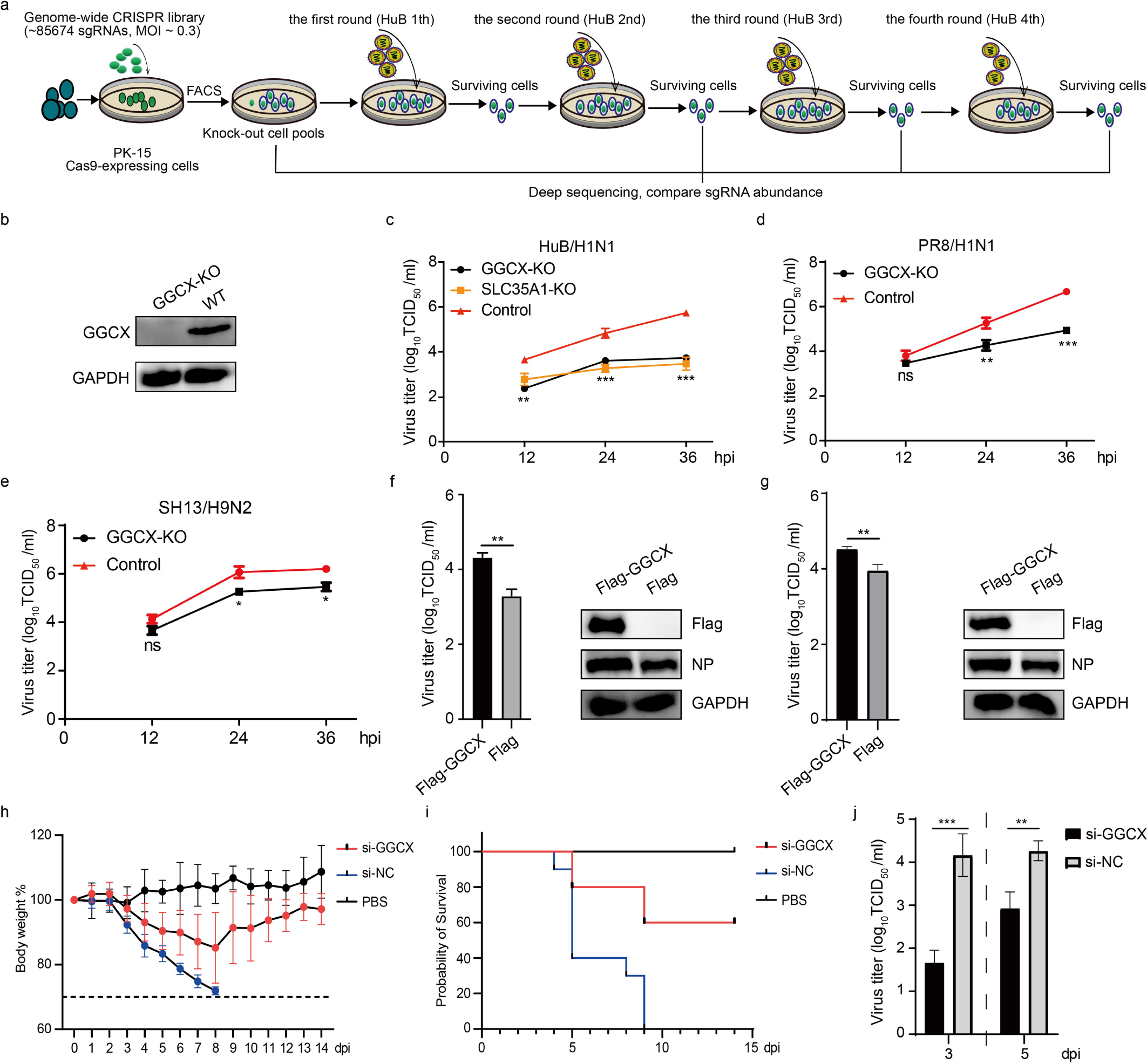
Genome-wide CRISPR screen identifies GGCX as EA H1N1 SIV host-dependent factor in porcine kidney cells. (a) A schematic diagram of the genome-wide CRISPR screening process in PK-15 cells. (b) Western blot analysis showing the expression of the GGCX protein in GGCX-KO and WT PK-15 cells, with GAPDH gene as an endogenous control. (c-e) Effects of GGCX-KO on the replication of influenza virus strains namely, (c) HuB/H1N1, (d) PR8/H1N1, and (e) SH13/H9N2. Viral titers were determined by TCID_50_. SLC35A1 is a IAV host-dependent factor and SLC35A1-KO cells were used as a positive control. (f-g) Restoration of GGCX promotes influenza virus replication. Exogenous GGCX (Flag-GGCX) and Flag negative control were transfected into (f) GGCX-KO PK-15 cells or (g) WT PK-15 cells, and viral titers determined by TCID_50_ and corresponding protein expression was detected by western blot. (h) Weight loss of HN/H1N1 strain infected mice after siRNA treatments. Mice with a body weight loss of more than 30% were euthanized according to the ethical principles of animal welfare. (i) Mortality of HN/H1N1 strain-infected mice after siRNA treatments. (j) Virus titers in the lungs of infected mice (n = 3) 3 days (left) and 5 days (right) after infection. (ns, *P* > 0.05; *, *P* < 0.05; **, *P* < 0.01; ***, *P* < 0.001).

To explore the role of host-mediated post-translational modifications (PTMs) in EA H1N1 SIV pathogenesis, an integral membrane protein, GGCX, was selected as it had been previously reported to catalyze the post-translational carboxylation of several proteins that convert specific peptide-bound glutamate (Glu) to γ-carboxyglutamate (Gla) ^25–27^. We employed CRISPR/Cas9 technology to successfully knock out the endogenous GGCX gene in PK-15 cells shown by the lack of GGCX protein expression and genomic base deletion (Fig. 1b and Extended Data Fig. 2a). GGCX deficiency did not appear to affect cell viability (Extended Data Fig. 2b). Interestingly, when compared with a well-known gene, SLC35A1, that is involved in the synthesis of sialic acid receptors and has been identified in many genome CRISPR screens ^24,28,29^, the ability of GGCX-knockout (KO) cells to inhibit HuB/H1N1 infectivity is the same as in SLC35A1-KO cells, suggesting that GGCX is key for efficient EA H1N1 SIV infection (Fig. 1c). We further evaluated the role of GGCX in the infection of other IAV strains and observed significantly reduced virus titers in GGCX-deficient cells infected with human (A/Puerto Rico/8/1934, PR8/H1N1) or avian-origin (A/chicken/Shanghai/SC197/2013, SH13/H9N2 IAV strains (Fig. 1d-e). Furthermore, the inhibitory effect of GGCX-deficient cells was more pronounced for EA H1N1 SIV (HuB/H1N1) than for human-origin (PR8/H1N1) and avian-origin (SH13/H1N1) strains (Fig. 1c-e). We then evaluated whether rescue or overexpression of GGCX could restore or promote viral infection. Our results show that overexpression of porcine GGCX in GGCX-deficient or WT cells can restore or promote HuB/H1N1 infection (Fig. 1f-g). Thus, the results showed that GGCX promoted the infection of EA H1N1 SIV *in vitro*.

To further investigate the role of GGCX in EA H1N1 SIV replication *in vivo*, chemically cholesterol-conjugated and 2’-OME-modified siRNA targeting GGCX (si-GGCX) and negative control siRNA (si-NC) were nasally instilled into 6-week-old female BALB/c mice, and the mice were challenged with A/Hunan/42443/2015 (HN/H1N1) (Extended Data Fig. 2c). The knockdown efficiency of the chemically modified siRNA in mice were assessed (Extended Data Fig. 2d). Mouse weight loss and survival were monitored daily for 14 days post-challenge. GGCX knockdown mice exhibited slightly attenuated infection as measured by reduced weight loss and increased survival rates compared to siRNA control mice (Fig. 1h-i). We also observed a significant decrease in viral titers in the lungs of GGCX knockdown mice compared to the control mice (Fig. 1j). Histopathological analysis of the infected control siRNA-treated mice’s lungs demonstrated moderate to severe bronchiolar necrosis, pulmonary oedema, and inflammatory cell infiltration (Extended Data Fig. 2e). Conversely, the examination of the infected GGCX knockdown mice’s lungs revealed a noteworthy reduction in the infiltration of lymphoid tissue compared to control mice (Extended Data Fig. 2e). Meanwhile, weaker viral nucleoprotein (NP) antigen signals were detected in the lungs of GGCX knockdown mice compared to control mice (Extended Data Fig. 2f). These results demonstrated that GGCX knockdown in the lungs of mice significantly contributed to a protective effect against EA H1N1 SIV challenge. Taken together, GGCX was identified as a porcine host-dependent factor for EA H1N1 SIV infection by an unbiased genome-wide CRISPR-Cas9 loss-of-function screen.

### GGCX catalyzes carboxylation modification of viral hemagglutinin

As previous reports established GGCX as an integral membrane protein catalyzing the post-translational carboxylation of several vitamin K-dependent (VKD) proteins ^25–27^, we hypothesized that GGCX participated in EA H1N1 SIV infection through post-translational carboxylation modification of viral proteins. To identify the viral protein targeted by GGCX-mediated carboxylation, we infected WT PK-15 cells with the HuB/H1N1 strain and conducted co-immunoprecipitation (Co-IP) experiments using anti-Gla antibodies. The results confirmed that the carboxylation modification of viral HA protein by immunoprecipitating carboxylated proteins (Fig. 2a). To validate the interaction between GGCX and HA, Co-IP experiments using anti-HA (Fig. 2b) or anti-GGCX (Fig. 2b) antibodies with WT cells infected with HuB/H1N1 strain confirmed the interaction between GGCX and viral HA proteins.

**Figure 2.**
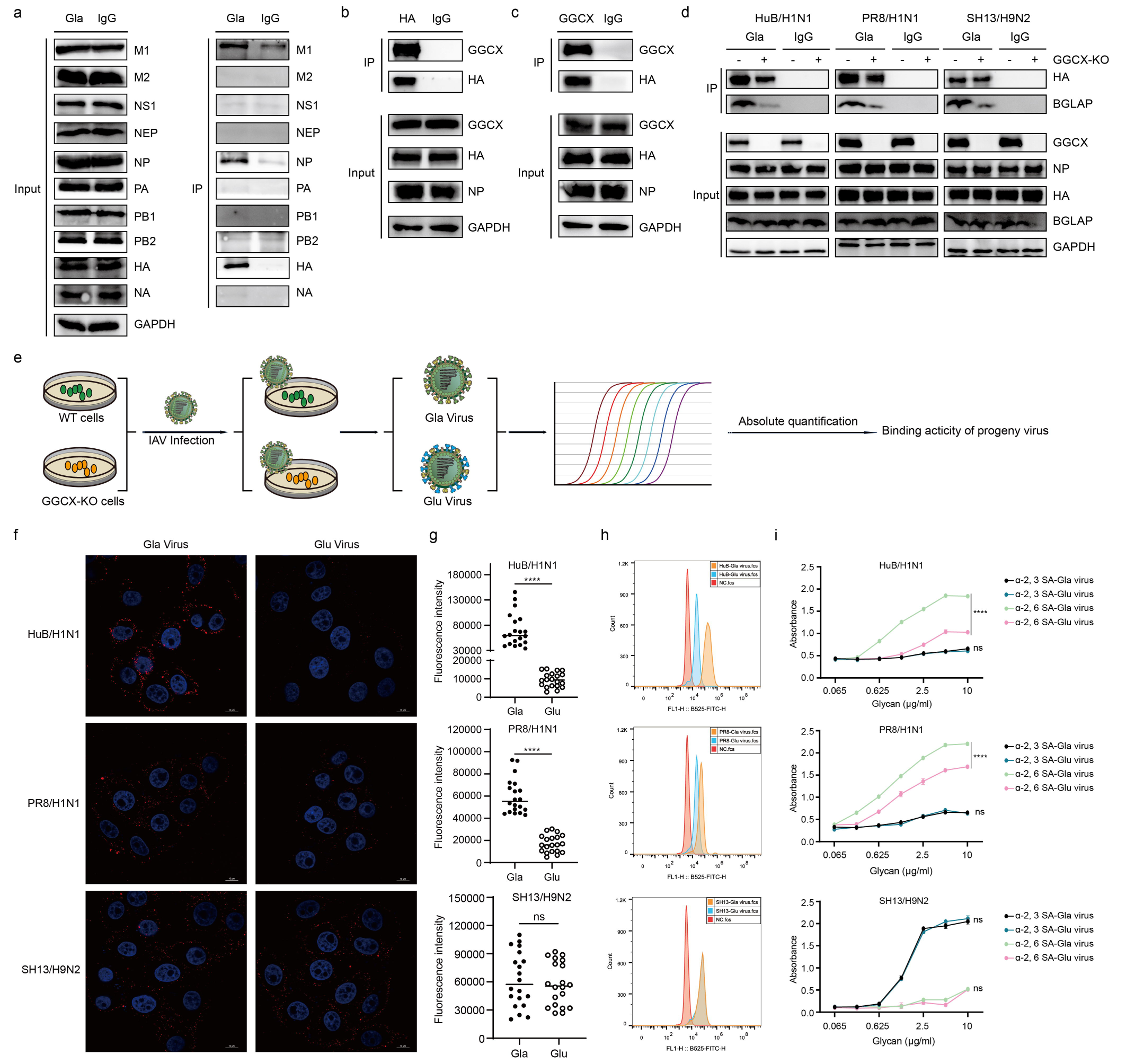
Hemagglutinin carboxylation modification by GGCX was required for progeny virus attachment. (a) Viral HA protein was modified for carboxylation. WT cells were infected with HuB/H1N1 strain, and cell lysates were prepared for Co-IP assays using anti-Gla or control IgG antibody, followed by western blot analysis to detect the co-precipitated viral proteins. (b-c) Interaction between endogenous GGCX and viral HA. WT cells were infected with HuB/H1N1 strain, and cell lysates were prepared for Co-IP assays using (b) anti-HA or (c) anti-GGCX with corresponding control IgG antibody, followed by western blot analysis to detect co-immunoprecipitated proteins. (d) Effects of GGCX-KO on different viral HA carboxylation modifications. GGCX-KO and WT cells were infected with three types of IAV strains. Cell lysates were prepared for Co-IP assays using anti-Gla or control IgG antibodies, followed by western blot analysis to detect co-precipitated proteins. BGLAP was used as a positive control. (e) Flowchart to determine the effects of GGCX-KO on progeny virus attachment. (f-h) Determination of binding activities of progeny Gla or Glu viruses. Equal amounts of progeny Gla or Glu viruses of HuB/H1N1, PR8/H1N1, and SH13/H9N2 were used to infect the WT PK-15 cells, respectively, followed by incubation with anti-influenza virus HA protein antibody. The HA proteins were then analyzed by (f) confocal microscopy (the red and blue fluorescence respectively indicated the HA protein and nucleus) and (g) average fluorescence intensity analysis as shown in (f) and (h) flow cytometry. Scale bars, 10 μm. (i) Sialic acid receptor binding preferences of progeny Gla or Glu virus. Equal amounts of progeny Gla or Glu viruses of HuB/H1N1, PR8/H1N1, and SH13/H9N2 were used to incubate with biotinylated sugar mimics of SA receptors, respectively, and ELISA assays were performed to determine receptor binding activities. (ns, *P* > 0.05; ****, *P* < 0.0001).

Interestingly, GGCX-KO resulted in reduced virus replication, with varying inhibitory effects observed among different IAV strains (Fig. 1c-e). We investigated whether this phenomenon was associated with the degree of carboxylation modification of viral protein mediated by GGCX. GGCX-KO and WT cells were infected with HuB/H1N1, PR8/H1N1, and SH13/H9N2, respectively, and carboxylation modified viral HA proteins were precipitated and detected by western blot assay. The results indicated the presence of carboxylated HA proteins in all three types of IAV-infected cells, with reduced expression in GGCX-KO cells, except in the SH13/H9N2-infected cells (Fig. 2d). Moreover, the carboxylation modification levels of HuB/H1N1 HA protein decreased more than that of PR8/H1N1, attributed to GGCX knockout (Fig. 2d). As the inhibitory effect of GGCX-deficient cells was more pronounced for EA H1N1 SIV (HuB/H1N1) than for human-origin PR8/H1N1 and avian-origin SH13/H9N2 (Fig. 1c-e), it was suggested that reducing viral infection was synchronous with the reduction of HA carboxylation modification for three IAV strains. These results collectively suggest that GGCX catalyzes the carboxylation modification of viral HA, potentially playing a critical role in EA H1N1 SIV infection.

### Carboxylation modification of viral HA by GGCX promotes receptor binding activity of progeny virus to host cells

GGCX promotes EA H1N1 SIV infection, and catalyzes the carboxylation modification of viral HA protein, which is involved in attachment to host cells. Therefore, carboxylation modification of HA may determine the receptor binding activity of progeny viral HA. To test our hypothesis, progeny viruses propagated in WT cells (Gla virus) or carboxylation modification insufficient GGCX-KO cells (Glu virus) were quantified by absolute quantitative real-time PCR, then the equivalent progeny Gla and Glu viruses were used to infect WT PK-15 cells and viral HA binding activities were visualized (Fig. 2e). We found that the progeny Glu viruses of HuB/H1N1 and PR8/H1N1 had a lower binding activity compared to the Gla viruses. In contrast, the progeny Gla and Glu viruses of SH13/H9N2 showed identical binding activities (Fig. 2f-g). Viral binding activities determined by flow cytometry were consistent with those obtained by confocal microscopy (Fig. 2h). Taken together, we found HA binding activity was directly regulated by GGCX-mediated carboxylation modification.

To assess which type of SA receptor binding activity of HA was affected in GGCX-KO cells, ELISA-based assays were used to investigate the receptor binding profiles of progeny Gla and Glu viruses. We observed that the progeny HuB/H1N1 and PR8/H1N1 viruses preferentially bound to α-2,6 SA receptors, and progeny Glu viruses of HuB/H1N1 and PR8/H1N1 had lower binding activity than the Gla viruses (Fig. 2i). In contrast, progeny SH13/H9N2 viruses preferentially bound to α-2,3 SA receptors and were not affected by GGCX-KO (Fig. 2i). Taken together, knocking out GGCX revealed different inhibitory effects on viral binding to different SA receptor types, and the binding activity of HA to the α-2, 6 SA receptors was regulated by GGCX-mediated carboxylation modification, indicating the critical role of GGCX in determining the receptor binding preferences of EA H1N1 SIV.

### GGCX-mediated carboxylation modification of viral HA 225E promotes its binding activity to **α**-2, 6 SA receptor

Since GGCX regulates the receptor binding activity of the progeny virus, we aimed to identify the carboxylation modification sites of viral HA catalyzed by GGCX. Thus, GGCX-KO and WT PK-15 cells were infected with the HuB/H1N1, and viral HA proteins were precipitated using anti-HA antibodies, followed by liquid chromatography-tandem mass spectrometry (LC-MS) analysis (Fig. 3a). The results revealed that, in addition to the 9 identical carboxylation modification sites (24, 106, 115, 118, 216, 246, 399, 408, and 427) identified in both HA proteins expressed in virus-infected GGCX-KO and WT cells, 13 distinct carboxylation modification sites (37, 97, 175, 225, 341, 387, 404, 433, 435, 450, 494, 501, and 502) were exclusively identified in HA proteins expressed in virus-infected WT cells (Supplementary Table 2). This suggests that these sites may play a crucial role in HA binding activity, which could be compromised by GGCX-KO (Fig. 3b). As GGCX-KO cells exhibited different effects on binding activity of progeny virus of EA H1N1 SIV (HuB/H1N1) and AIV (SH13/H9N2), the conservation of the 13 carboxylation sites was analyzed among the EA H1N1 SIV and different AIV subtypes. Six glutamic acid (E) carboxylation sites (97, 175, 225, 387, 450, and 494) were selected for further study, due to the glycine (G) or aspartic acid (D) substitutions at the corresponding carboxylation sites in AIV subtypes (Fig. 3c and Extended Data Fig. 3).

**Figure 3.**
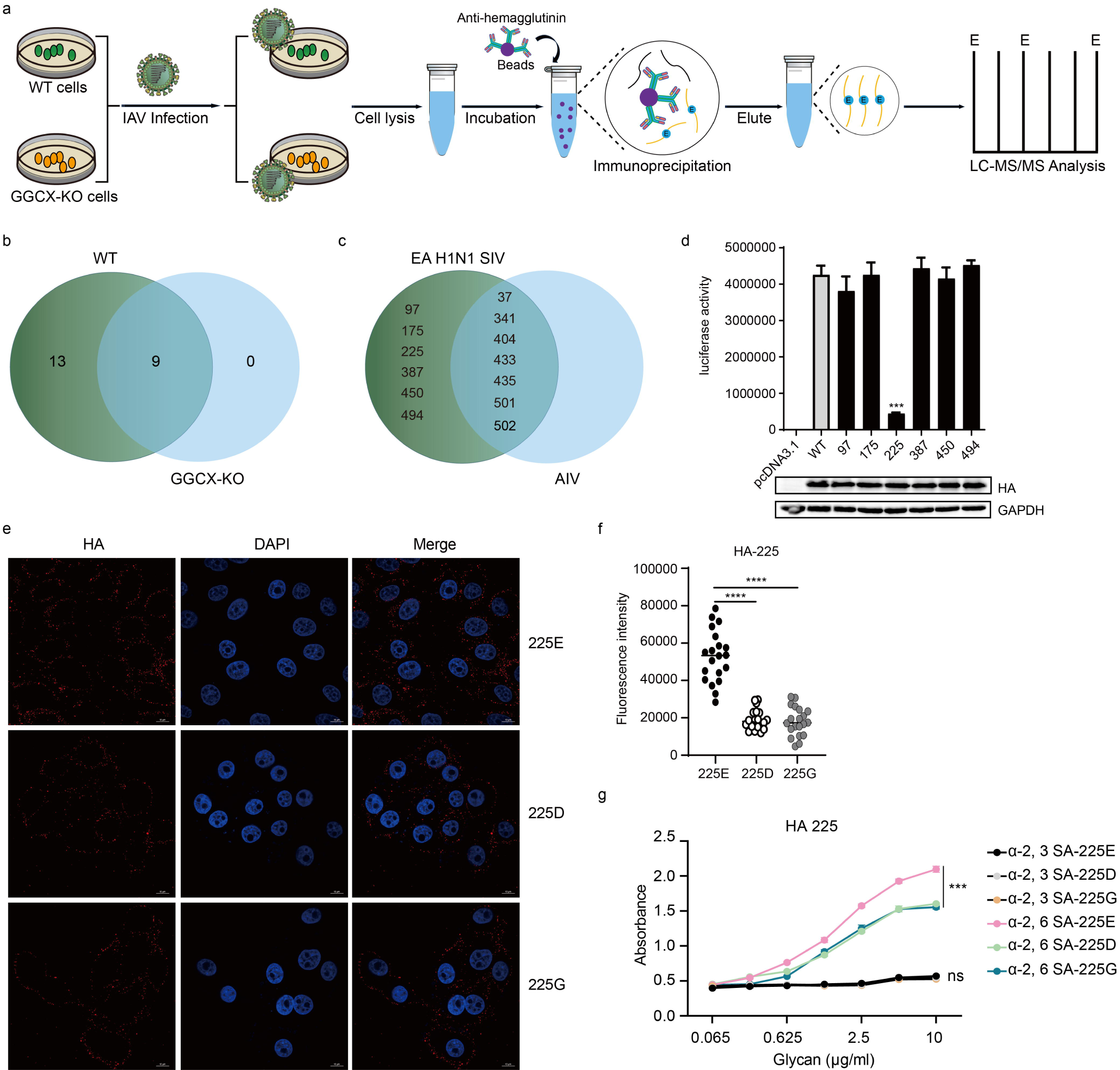
GGCX-mediated Carboxylation modification of viral HA 225E promotes its binding activity to. α**-2, 6 SA receptors.** (a) Flowchart showing identification of carboxylation modification sites of viral HA by LC-MS. (b) Venn diagram of identified carboxylation modification sites in viral HA proteins expressed in infected GGCX KO and WT cells. (c) The Venn diagram analysis of the conservation of glutamic acid (E) at the carboxylation modification sites in EA H1N1 SIV and different AIV subtypes. (d) Identification of key carboxylation modification sites of viral HA. HEK293T cells were transfected with psPAX2, pLenti-luc, and WT or mutant viral HA plasmids to generate pseudoviruses. The pseudoviruses were then used to infect WT PK-15 cells and luciferase assays were performed. Protein expression levels were evaluated by western blot. (e-f) Binding activities of HA 225E and mutant viruses. Equal amounts of HA 225E, 225D, and 225G viruses were each used to infect the WT PK-15 cells and incubated on ice for 1 hour, followed by incubation with an anti-influenza virus HA protein antibody. Then, the HA proteins were analyzed by (e) confocal microscopy and (f) average fluorescence intensity analysis as shown in (e). Scale bars, 10 μm. (g) Sialic acid receptor binding preferences of HA 225E and mutant viruses. Equal amounts of HA 225E, 225D, and 225G viruses were used to incubate biotinylated sugar mimics of sialic acid receptors, and ELISA assays were performed to determine the receptor binding activities. (ns, *P* >0.05; ***, *P* < 0.001; ****, *P* < 0.0001).

Carboxylation site mutant and WT pseudoviruses were generated, and their binding activities were evaluated through a luciferase assay. The results indicated that the mutant HA E225A inhibited the binding activities of pseudoviruses (Fig. 3d), highlighting the critical role of the carboxylation modification of HA 225E in determining virus binding activities.

To explore the impact of carboxylation modification of HA 225E on receptor binding activity of EA H1N1 SIV, we introduced HA E225D or E225G substitutions into the HN/H1N1 virus, generating two corresponding mutant viruses. Equal amounts of mutant and WT viruses were then used to infect PK-15 cells, and the binding activities were compared through confocal microscopy. The results demonstrated that mutant viruses with HA 225D and 225G exhibited lower binding activities than the WT HA 225E virus (Fig. 3e-f). Additionally, we observed that the binding activities of the two mutant viruses to α-2,6 SA receptor were decreased compared to the WT virus, while binding activities to α-2,3 SA receptor showed no differences between the different virus types (Fig. 3g). Collectively, these results indicate that GGCX-mediated 225E carboxylation modification can regulate the binding activities of viral HA to α-2,6 SA receptor.

### GGCX mediates the HA 225E substitution of EA H1N1 SIV

The EA H1N1 SIVs originated from Eurasian avian H1N1, and the viral HA 225E site has been reported to be involved in HA receptor binding sites and determines the switch of EA H1N1 SIV receptor binding specificity from α-2,3 SA receptor to α-2,6 SA receptor ^30,31^. When analyzing the conservation of HA 225E in H1N1 AIV and EA H1N1 SIV, we found that HA 225E was more conserved in EA H1N1 SIV than in H1N1 AIV (Fig. 4a). Furthermore, the conservatism of HA 225E in EA H1N1 SIV increased over time while that of HA 225G decreased significantly (Fig. 4b). This suggests that HA 225E plays a key role in determining the evolutionary adaptation of EA H1N1 SIV.

**Figure 4.**
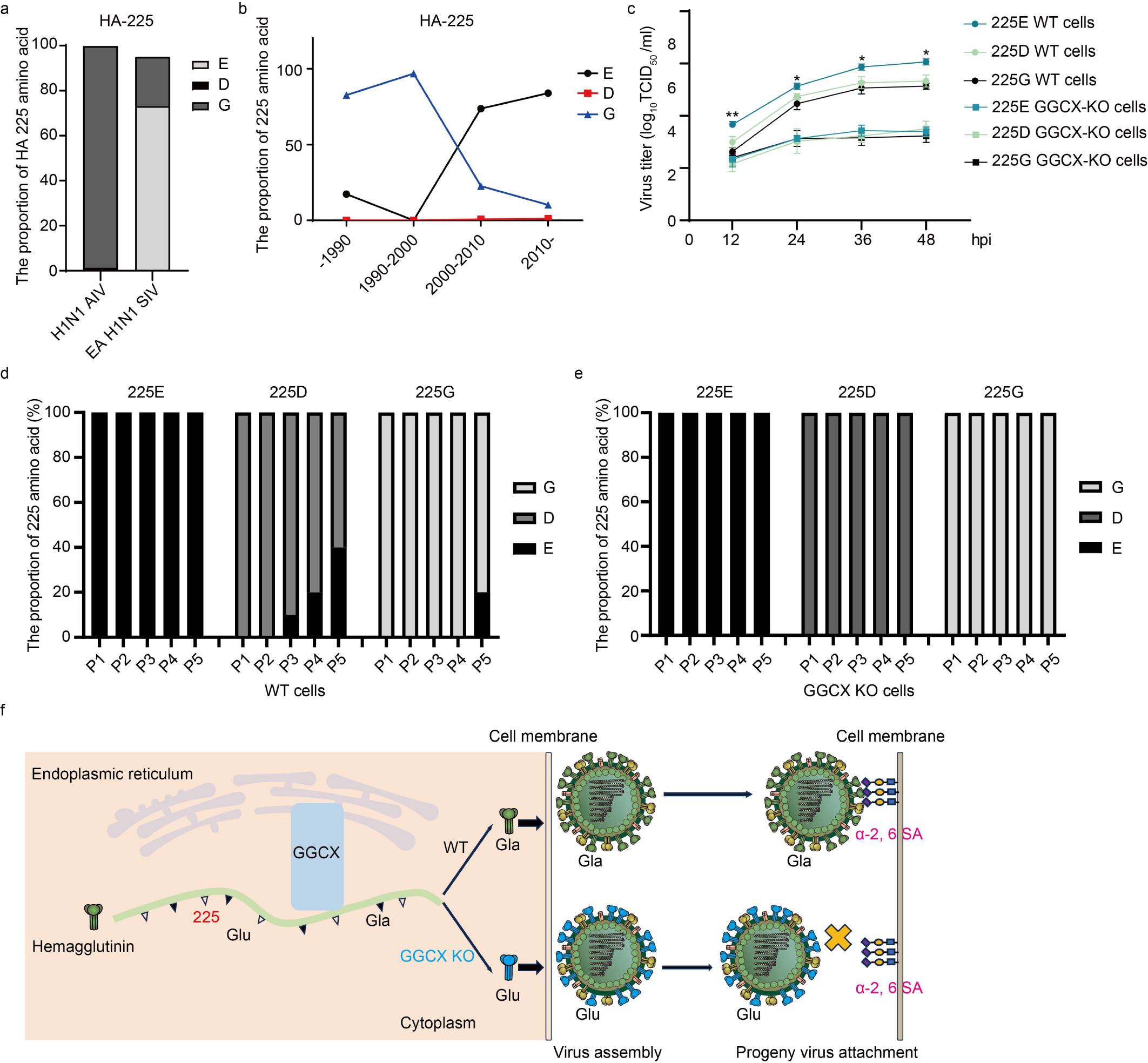
GGCX catalyzed the substitution adaption of EA H1N1 SIV HA 225E. (a) Conservation analysis of the HA 225E site in H1N1 AIV and EA H1N1 SIV. (b) Conservation change analysis of the EA H1N1 SIV HA 225E site over time. (c) Virus growth kinetics curves of HA 225E and mutant viruses in WT and GGCX-KO PK-15 cells. HA 225E, 225D, and 225G viruses were used to infect WT and GGCX-KO PK-15 cells, respectively, at MOI=0.1, and the virus titers at the indicated time points were determined by TCID_50_. (*, *P* < 0.05; **, *P* < 0.01) (d-e) EA H1N1 SIV passage and sequencing. HA 225E, 225D, and 225G viruses were used to infect the (d) WT and (e) GGCX-KO PK-15 cells, respectively, at MOI=0.1 for 48 hours, the progeny viruses were collected and sequenced to infect the WT and GGCX-KO cells, respectively, and the viruses from each round were subjected to sequencing. (f) Working model for the regulation of GGCX in the receptor binding activity of progeny EA H1N1 SIV.

Since the GGCX-catalyzed carboxylation of HA 225E determines the receptor binding activity of EA H1N1 SIV to the α-2,6 SA receptor, GGCX might be responsible for the evolutionary adaptation of EA H1N1 SIV HA G225E. To further clarify the role of GGCX in the evolutionary adaptation of EA H1N1 SIV HA 225E, WT HA 225E and mutant viruses were infected into GGCX-KO and WT cells, respectively, and viral growth curves were documented. The result showed that the titers of the proliferating HA 225E virus from the WT PK-15 cells were significantly higher than those of the mutant virus. However, the titer of the virus proliferating from the GGCX-KO cells was almost the same as that of the mutant virus (Fig. 4c).

Meanwhile, the progeny viruses infecting GGCX-KO and WT cells were sequenced sequentially after serial passage. It was found that the HA 225D and HA 225G mutant viruses proliferating from WT PK-15 cells gradually reverted to HA 225E, whereas the viruses proliferating from GGCX-KO cells remained unchanged (Fig. 4d-e), suggesting that the substitution of HA 225E was catalyzed by GGCX. Taken together, GGCX is a host factor required for HA 225E carboxylation modification and mediates the HA 225E substitution of EA H1N1 SIV.

## Discussion

Over an extended evolutionary timeframe, EA H1N1 SIVs have gradually accumulated increased affinity for binding to α-2,6 SA receptor, facilitated in part by the D225E substitution in the viral HA. This substitution, has the potential to contribute to the emergence of the next influenza pandemic. However, the precise factor driving this evolutionary substitution has poorly understood. Our investigation identified that GGCX promotes EA H1N1 SIV infection through a genome-scale CRISPR screen conducted in porcine kidney cells. The research uncovered that GGCX-mediated post-translational carboxylation modification of viral HA played a critical role in enabling progeny EA H1N1 SIV to bind to α-2, 6 SA receptor (Fig. 4e). Moreover, GGCX emerged as the determinant for the D225E substitution, actively promoting the evolutionary adaptation of EA H1N1 SIV. Collectively, GGCX plays a pivotal role in catalyzing the carboxylation modification of viral HA 225E, thereby fostering the receptor binding preferences of EA H1N1 SIV to the α-2, 6 SA receptor and promoting the evolutionary adaption of EA H1N1 SIV.

In contrast to previous genome-scale CRISPR screens primarily focused on the isolated human or avian influenza virus in their hosts, our study initiated a genome-scale CRISPR screen in porcine kidney cells to identify the host-dependency factors necessary for EA H1N1 SIV infection ^24,28,32^. Shared hits with other studies highlighted the enrichment of host factors involved in SA receptor biosynthesis and related glycosylation pathways, including SLC35A1, CMAS, ST3GAL4, and ALG5, emphasizing the critical role of SA receptor biosynthesis in multiple IAV strains ^28,33^. Additionally, unique genes specific to the EA H1N1 SIV strain (HuB/H1N1) and its corresponding host were identified, indicating their specific involvement in EA H1N1 SIV infection. GGCX was identified as essential for EA H1N1 SIV infection and was confirmed to be responsible for the HA 225E substitution through its previously unreported carboxylation modification function on the viral HA. Similar to other post-translational modifications such as glycosylation, methylation, and acetylation ^34–36^, carboxylation modification of viral HA can alter the structure and functions of the target HA protein, influencing the formation of the salt bridge structure between amino acids D225/E225 and K222, as well as virion assembly and budding efficiency ^19,20^. This may explain the switch in the SA receptor binding preferences of the EA H1N1 SIV from α-2,3 SA to the α-2,6 SA receptor. In addition, the protein structures of the viral HA protein, with and without carboxylation modification, would be analyzed to clarify the mechanism by which HA carboxylation modification alters the SA receptor binding preference and determining the evolutionary adaptation of EA H1N1 SIV.

Our study showed that the carboxylation modification insufficient progeny Glu virus had lower α-2,6 SA receptor binding activities than the Gla virus. Carboxylated modification of HA 225E was identified as a key regulator of progeny virus binding activities. However, the decreased binding activity observed in the 225E mutant was not sufficient to compensate for the KO effect of GGCX, suggesting the presence of other unidentified carboxylation modification sites. GGCX plays a critical role in the vitamin K cycle, which facilitates the γ-carboxylation and recycling of VK via GGCX and vitamin K epoxide reductase (VKOR), respectively ^25–27^. Therefore, carboxylation modification of viral HA may undergo dynamic cyclic changes, with carboxylation modification sites varying at different stages of viral HA function. To investigate additional functional carboxylation modification sites of HA, HA proteins expressed in infected cells at different post-infection timepoints are collected and subjected to LC-MS analysis. In addition, potential carboxylation modification sites located in the receptor binding domain of HA will be validated by structural analysis. Overall, the identification of functional carboxylation modification sites in HA highlights the importance of viruses with substitutions at these sites, which may have a higher propensity to bind to α-2,6 SA receptor and pose a potential threat to human health.

GGCX is required for infection of several IAV strains, including EA H1N1 SIV and AIV. Interestingly, carboxylation modification of the AIV (SH13) HA protein and progeny virus binding activities were not affected by GGCX-KO, suggesting that GGCX-regulated IAV infection involves alternative mechanisms. This observation suggests that carboxylation modification of other Gla proteins catalyzed by GGCX may also contribute to IAV replication. In addition to the HA protein, the viral M1 and NP proteins were also found to undergo carboxylation modification with potential regulation by GGCX, indicating a universal effect on IAV replication ^37,38^. Furthermore, host proteins that undergo carboxylation modification, such as coagulation factors, osteocalcin and matrix Gla proteins, have been implicated in viral protein cleavage, autophagy and immune pathways ^39–41^, providing an alternative explanation for the inhibition of IAV replication. In conclusion, our study highlights the multifaceted role of GGCX in the regulation of IAV replication and suggests that it may serve as a potential target for the development of IAV therapeutics.

In conclusion, we used CRISPR/Cas9-based high-throughput loss-of-function screening to identify cellular factors involved in EA H1N1 SIV infection. Next, we found that GGCX catalyzes the carboxylation modification of the viral HA protein. This modification is essential to promote the binding capacity of progeny EA H1N1 SIV to α-2, 6 SA receptor. In particular, the GGCX-catalyzed carboxylation modification was found to be responsible for the substitution of HA 225E during the evolutionary adaptation of EA H1N1 SIV. These findings provide valuable insights into the mechanisms underlying the receptor binding preferences adaptation and the potential evolutionary adaptation of EA H1N1 SIV. They have important implications for the development of interventions aimed at disrupting transmission pathways and mitigating the risk of future influenza pandemics.

## Methods

### Ethics statement

Approval for the animal experiments carried out in this study was obtained from the Committee on the Ethics of Animal Experiments at Huazhong Agricultural University (No. HZAUMO-2023-0286).

### Cells

Human embryonic kidney 293T cells (HEK293T, Cat# CRL-3216), Madin-Darby canine kidney (MDCK, Cat# CCL-34) cells, and Porcine Kidney-15 (PK-15, Cat# CCL-33) cells were purchased from the American Type Culture Collection (ATCC, Manassas, VA, USA). Stably expressing Cas9 PK-15 (PK-15-Cas9) cells were established through puromycin screening. All cell lines were cultured at 37°C in a 5% CO_2_ humidified atmosphere using RPMI 1640 (SH30809.01, HyClone, USA) or Dulbecco’s modified Eagle’s medium (DMEM) (SH30243.01, HyClone, USA) supplemented with 10% fetal bovine serum (FBS) (FSP500, ExCell, China).

### Viruses and reverse genetics

The IAVs used in this study were A/Swine/HuBei/221/2016 (HuB/H1N1), A/Puerto Rico/8/1934 (PR8/H1N1), A/chicken/Shanghai/SC197/2013 (SH13/H9N2), and A/Hunan/42443/2015 (HN/H1N1). Recombinant viruses were generated in the genetic background of A/Hunan/42443/2015 (HN/H1N1) using an eight-plasmid reverse genetic system as described previously ^42^. All other viruses were amplified using 10-day-old embryonic chicken eggs and titrated by determining TCID_50_ values on MDCK cells. All experiments with A/chicken/Shanghai/SC197/2013 (SH13, H9N2) virus were performed in an animal biosafety level 3 laboratory at Huazhong Agricultural University.

### Plasmids

Lentiviruses was produced using the lenti-sgRNA-EGFP vector, along with the pMD2.G and psPAX2 plasmids. Pseudoviruses were generated using the pLenti-luc, a generous contribution from Dr Rui Luo of Huazhong Agricultural University. For the construction of the lentiviral sgRNA vector, paired sgRNA oligonucleotides (50 μM per oligo) were annealed and cloned into lenti-sgRNA-EGFP vector, which was linearized with *Bbs*I (R3539, NEB, USA). The p3xFlag-GGCX (Flag-GGCX) was constructed by cloning the full-length cDNA, amplified by PCR, cloned into the p3×Flag (Flag) vector digested with *Hin*dIII/*Xba*I. Eight segments of A/Hunan/42443/2015 (HN/H1N1) were inserted into the pHW2000 vector, and mutant HA genes targeting amid acid 225 were generated by PCR-based site-directed mutagenesis, confirmed by sequencing.

### Antibodies and reagents

The antibodies and reagents used in the study were as follows: Rabbit anti-GGCX (16209-1-AP, Proteintech, China); mouse anti-Gla (3570, Biomedica, Canada), mouse anti-Flag tag (F1804, Sigma-Aldrich, USA); rabbit anti-BGLAP (A6205, ABclonal, China); mouse anti-glyceraldehyde-3-phosphate dehydrogenase (GAPDH) (CB100127, California Bioscience, USA); rabbit anti-IAV NP and HA (GTX125989 and GTX127357, GeneTex, USA); horseradish peroxidase-conjugated anti-mouse and anti-rabbit (BF03001 and BF03008, Beijing Biodragon Immunotechnologies, China); goat anti-Cy3 anti-rabbit IgG (H+L) (AS007, ABclonal, China), and 4′,6′-diamidino-2-phenylindole (1:5,000) (C1002, Beyotime, China).

### Genome-scale CRISPR screening in pig

To conduct CRISPR screening, approximately 6×10^7^ genome-scale PK-15 mutant cell libraries underwent infection with A/Swine/HuBei/221/2016 (HuB/H1N1) at an MOI of 0.01 in DMEM devoid of FBS. The cells were then incubated at 37 [and 5% CO_2_. Following a 1.5-hour incubation, the initial inoculum was replaced with fresh DMEM supplemented with 2.5% BSA (Cat# A4161, Sigma-Aldrich), 0.25μg/ml TPCK (Cat# 4370285, Sigma-Aldrich), and 1% penicillin-streptomycin (Cat# P4333, Sigma-Aldrich). Surviving cells were collected 10 days post-infection and expanded for subsequent rounds of infection. After four rounds of screening, high throughput sequencing was applied to the surviving cells from the second, third, and fourth rounds of challenge, followed by the analysis of candidate genes.

### Generation of GGCX knockout cell line using CRISPR/Cas9

Individual sgRNA constructs targeting GGCX were generated and incorporated into the lenti-sgRNA-EGFP vector. Lentiviruses were produced following established protocols ^43^. These lentiviruses were then transduced into PK-15-Cas9 cells. Transduced cells were selected using fluorescence-activated cell sorting (FACS). Monoclonal cells were obtained through the limiting dilution method and subsequently expanded. Confirmation of GGCX-KO cells was achieved through Sanger sequencing and western blot analysis.

### Cell viability assay

To examine the impact of GGCX-KO on cellular proliferation, the viability of GGCX-KO cells and WT cells was measured through CCK-8 activity, following the manufacturer’s instructions ^43^. Briefly, cells were seeded onto 96-well plates, and their viabilities were measured at 12-, 24-, and 36-hours post-seeding. CCK-8 reagent (CK04-500T, Dojindo Molecular Technologies, Japan) was applied to each well. and the subsequent measurement of absorbance at 450 nm was conducted using a microplate reader after a 1-hour incubation at 37°C in dark.

### Virus infection and titration

To evaluate the impact of GGCX-KO on viral replication, negative control and GGCX-KO cells were independently seeded in triplicate within 12-well plates. For influenza A virus (IAV) infection, cells underwent two times washes with DMEM, followed by incubation with diluted virus at the MOI of 0.01 for 1 hour. Subsequently, cells were again washed twice with DMEM and replenished with fresh infection medium (DMEM supplemented with 0.2 μg/ml TPCK-treated trypsin (T1426, Sigma-Aldrich, USA)). Supernatants were collected at designated time points post-infection, and viral supernatants were serially diluted with DMEM. Eight replicates of each dilution were added to the wells, and the 50% tissue culture infectious dose (TCID_50_) was calculated using the Reed-Muench method 72 hours after infection ^44^.

### Mouse models for GGCX knockdown

For an in-depth exploration of GGCX’s role in EA H1N1 SIV infection *in vivo*, we synthesized cholesterol-conjugated and 2’-OME-modified si-GGCX or si-NC (GenePharma, China) and administered them nasally to 6-week-old BALB/c female SPF mice on days 0 and 2 ^45^. Subsequently, the mice were either challenged with 30 pfu of HN/H1N1 or mock infected on day 1. Daily monitoring of body weight loss and survival occurred over 2 weeks post-infection (n = 10). Mice exceeding a 30% loss in initial body weight were humanely euthanized. On 3 and 5 days post-challenge, a subset of mice from each group (n = 3) underwent anesthesia, and sacrifice and their lungs were either homogenized and/or fixed in 4% formaldehyde. The homogenized lung samples were utilized for assessing gene expression as well as virus titers. The fixed mouse lung samples were used for hematoxylin & eosin (H&E) and immunofluorescence staining for histopathological analysis.

### Western blot and immunoprecipitation

For immunoprecipitation, GGCX-KO and WT cells were infected with the IAV strains at 0.01 MOI for 12 hours. Cells were washed with cold phosphate-buffered saline (PBS) and lysed with NP-40 lysis buffer (P0013F, Beyotime, China) containing protease inhibitor cocktail (04693132001, Roche, Switzerland). Cell lysates were incubated overnight at 4 [with Dynabeads (Sc-2003, Santa Cruz Biotechnology, USA) conjugated with antibodies against either the γ-carboxyglutamyl (Gla) residues or control IgG antibodies. Protein-antibody-Dynabeads complexes were washed three times with NP-40 lysis buffer and analyzed by western blot.

### Absolute quantitative real-time PCR

Viral RNA was extracted from cell suspensions using TRIzol Regent (15596018, Invitrogen, USA) according to the manufacturer’s protocol. Extracted viral RNAs were used as a template to generate cDNA using reverse transcriptase (RK20403, ABclonal, China). Quantitative real-time PCR (qRT-PCR) (ABI Vii7A, USA) was performed using SYBR GREEN (RK21203, ABclonal, China). The constructed plasmid expressing full length of viral NP (pcDNA3.1-NP) was used as a standard to generate a standard curve. The amount of viral RNA was calculated according to the formula provided by the standard curve^46^.

### Receptor binding activity assay

Serial dilutions of Neu5Acα2-3Galβ1-4Glcβ-sp4-PAA-biot (0060-BP, GlycoNZ, New Zealand) and Neu5Acα2-6Galβ1-4GlcNAcβ-sp3 (0997-BP, GlycoNZ, New Zealand) were applied to pre-streptavidin-coated high-capacity plates (15500, Thermo Scientific, USA) and incubated at 4 [for overnight ^47^. Subsequently, the plates were washed three times with PBS, followed by incubation with 2% PBSA for 1 hour at room temperature and three additional PBS washes. Diluted influenza virus was then added to the plates and allowed to incubate at 4 [overnight. After five washes with PBST, the plates were incubated with chicken anti-influenza virus serum for 4 hours at 4[, washed with PBST, and incubated with HRP rabbit anti-chicken (IgG) (H+L) (AS030, ABclonal, USA). TMB substrate (CW0050S, CWBIO, China) was added to the plates to react for 20 minutes at room temperature in dark and stopped with 0.5 M H_2_SO_4_. OD values were recorded at 450 nm wavelength using a multimode reader (EnVision).

### HA binding assays

Cell surface binding of HA was performed with quantified progeny Gla and Glu virus. Briefly, wild-type PK-15 cells were incubated with progeny Gla and Glu virus of HuB/H1N1, PR8/H1N1, and SH13/H9N2 at an MOI of 5 for 1 hour on ice and unbound protein was washed with PBS. Cells were then fixed with 4% formaldehyde for 10 minutes and incubated with 1% (wt/vol) bovine serum albumin (BSA) for 1 hour at room temperature. The amount of bound viral HA was measured by cell surface staining for viral HA with a polyclonal anti-HA antibody followed by a second antibody conjugated to anti-Cy3 goat anti-rabbit IgG (H+L) (AS007, ABclonal, China) and observed by confocal microscopy analysis.

### Flow cytometry

Wild-type PK-15 cells were infected with titrated progeny Gla and Glu virus of HuB/H1N1, PR8/H1N1, and SH13/H9N2 at an MOI of 5 and incubated at 4 [for 1 hour. Subsequently, the cells were collected, fixed with 4% paraformaldehyde for 10 minutes, and washed three times with PBS. After a 2-hour incubation with 1% PBSA, the cells were incubated with anti-influenza virus HA (GTX127357, GeneTex, USA) for additional 2 hours. Staining of the cells was achieved using FITC-conjugated goat anti-rabbit IgG (H+L) (5230-0359, SeraCare, USA), and positive cells were analyzed through Cytoflex-LX.

### Liquid chromatography-tandem mass spectrometry

GGCX-KO and WT cells were infected with the HuB/H1N1 strain virus at MOI of for 9 hours. Subsequently, cells were washed with cold PBS and lysed using NP-40 lysis buffer (P0013F, Beyotime, China) supplemented with a protease inhibitor cocktail (04693132001, Roche, Switzerland). The resulting cell lysates were each incubated overnight at 4 [with Dynabeads (Sc-2003, Santa Cruz Biotechnology, USA) pre-conjugated with antibodies against the viral HA protein (GTX127357, GeneTex, USA). Complexes of protein-antibody-Dynabeads were washed three times with NP-40 lysis buffer and then subjected to SDS-PAGE. The gel bands corresponding to the molecular weight of viral HA were collected and digested by chymotrypsin and Trypsin & ASP-N, subsequently the digested proteins were analyzed by LC-MS (Biotechpack, China). The glutamic acid (E) sites which added a corresponding carboxyl molecular weight (44 Da) were identified as the carboxylation modification sites.

### Pseudoviruses packaged and assayed for luciferase activity

Pseudoviruses were produced by co-transfecting HEK293T cells with psPAX2, pLenti-luc and plasmids encoding either the WT or the carboxylation site mutant (97, 175, 225, 387, 450 and 494 E/A) viral HA, using Lipofectamine™ 2000 (11668019, Invitrogen, USA) ^48^. Supernatants were harvested at 60- and 72-hours post-transfection and filtered through a 0.45 μm filter. Wild-type PK-15 cells were seeded (cell density of 20%) in 12-well plates and incubated in 1 mL media containing pseudoviruses for transduction. Following a 1-hour incubation on ice, the transduced cells were replenished with fresh media, lysed, and the luciferase activities of the pseudoviruses were gauged using a luciferase assay system (E1501, Promega, USA).

### Virus passage and sequencing

GGCX-KO and WT cells were infected with HA 225E, 225D, and 225G HN/H1N1 viruses for 48 hours. The supernatants containing the progeny viruses were collected, and these viruses were subjected to sequential rounds of infection on GGCX-KO and WT cells for an additional four cycles each. Viral RNAs obtained from these five rounds were extracted using TRIzol Regent (15596018, Invitrogen, USA). The extracted viral RNAs served as templates for cDNA synthesis using reverse transcriptase (RK20403, ABclonal, China). Viral HA segments were PCR-amplified and cloned into pMD-18T vector (6011, TAKARA Beijing, China). Subsequently, 10 bacterial colonies were randomly selected for sequencing (Tsingke, China), and the proportions of HA amino acids 225E, 225D, and 225G were analyzed.

### Statistical analysis

All measurements were taken in triplicate, and the presented data are outcomes from at least two separate experiments. The results are shown as the mean ± standard deviation of the triplicate determinations. Statistical significance was ascertained by computing *P* values using the paired two-tailed Student’s t-test (ns, *P* >0.05; *, *P* < 0.05; **, *P* < 0.01; ***, *P* < 0.001; ****, *P* < 0.0001).

## Acknowledgments

We acknowledge the support of the National Key Laboratory of Agricultural Microbiology Core Facility for their aid in confocal microscopy and flow cytometry. Special thanks go to Xiao Xiao from Huazhong Agricultural University, China, for the meticulous review of the manuscript.

This work was supported by the National Key Research and Development Program (2021YFD1800204), the National Natural Science Foundation of China (32025036), the Fundamental Research Fund for the Central Universities (2662023PY005), Hubei Hongshan Laboratory (2022hszd005), the earmarked fund for CARS-41, and the Natural Science Foundation of Hubei Province (2021CFA016).

## Author contributions

H.Z., S.X., and W.B. conceived the project; J.Z., M.J., R.X., H.S., H.L., S.T., T.C., J.G., and Y.Z. conducted the experiments; J.Z., M.J., R.X., H.S., T.P., S.X., W.B., and H.Z. analyzed the data; J.Z., T.P., S.X., and H.Z. wrote and revised the paper. All authors reviewed and approved the final manuscript.

## Data availability

All data are available in the Article and its Supplementary Information. Source data are provided with this paper.

## Competing interests

The authors declare no conflict of interest.

Extended data and supplementary information are available for this paper. Correspondence and requests for materials should be addressed to H.Z.; S.X.; or W.B.

---

**Extended Data Figure 1.**
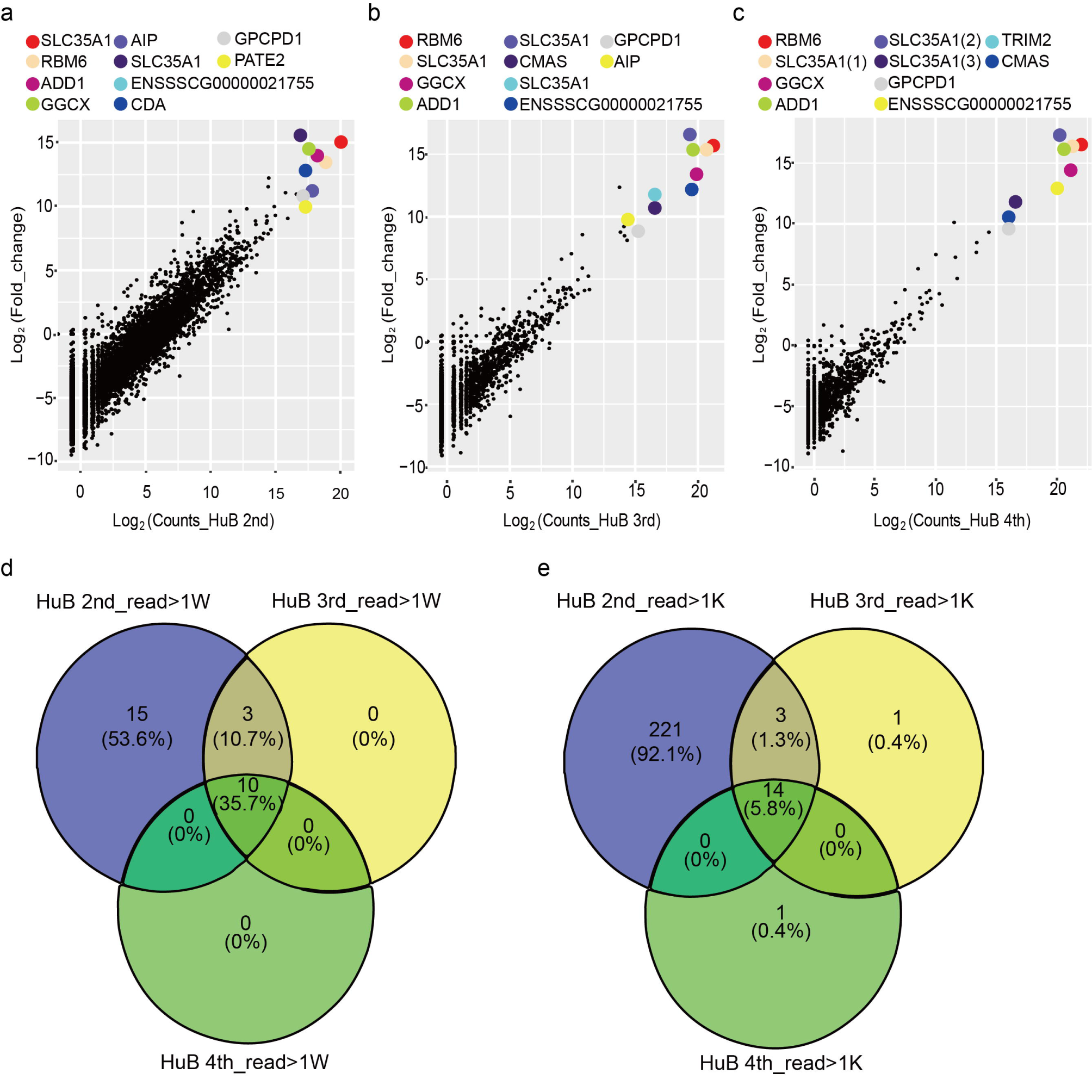
Analyzing of the sequenced sgRNA of EA H1N1 SIV screens after challenge. (a-c) Scatter plots comparing sgRNA targeting sequences frequencies and extent of enrichment vs the non-inoculated control mutant cell pool for the (a) second, (b) third, and (c) fourth rounds of EA H1N1 SIV screens after challenge. (d-e) Venn diagrams showing the overlapping enrichment of specific sgRNAs targeting sequences in the second, third, and fourth rounds of EA H1N1 SIV screens after challenge. For (d), among the reads over 10,000 for the sgRNAs; for (e), among the reads over 1,000 for the sgRNAs.

**Extended Data Figure 2.**
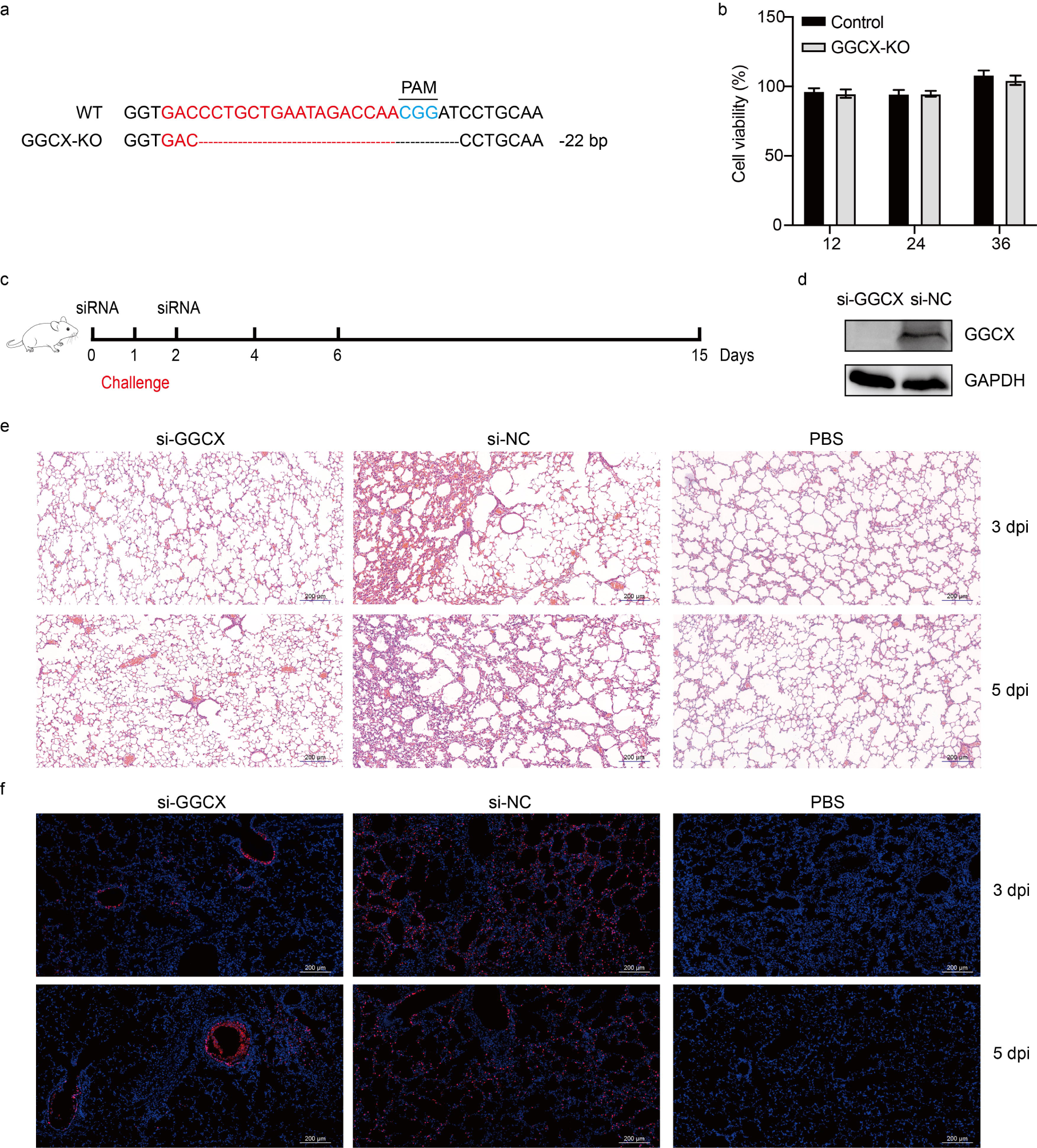
Determination of the knockdown efficiency of GGCX and its effect on EA H1N1 SIV infection after knockdown. (a) Sanger sequencing confirmation for the generated GGCX-KO cell lines using CRISPR/Cas9 technology. The sgRNA sequence was highlighted in red and NGG sequence was highlighted in blue. PAM: Protospacer Adjacent Motif. (b) Cell viability in GGCX-KO and WT PK-15 cells was determined using CCK-8 reagents over 36 hours. (c) Schematic of siRNA treatment and HN/H1N1 strain challenge in an experimental mouse model (n = 10). Mice were treated with siRNA the day before and after the viral challenge, and monitored for 14 days. (d) Western blot analysis of GGCX protein expression in GGCX-siRNA treated mice. (e) Hematoxylin and eosin (H&E) staining of pathological lesions in the lungs of GGCX knockdown mice infected with HN/H1N1 strain at 3 and 5 days post-challenge. Scale bars, 200 μm. (f) Immunofluorescence staining of lung sections from GGCX knockdown mice infected with HN/H1N1 strain at 3 and 5 days post-challenge. The viral NP antigen was stained red, and the nucleus was stained blue. Scale bars, 200 μm.

**Extended Data Figure 3.**
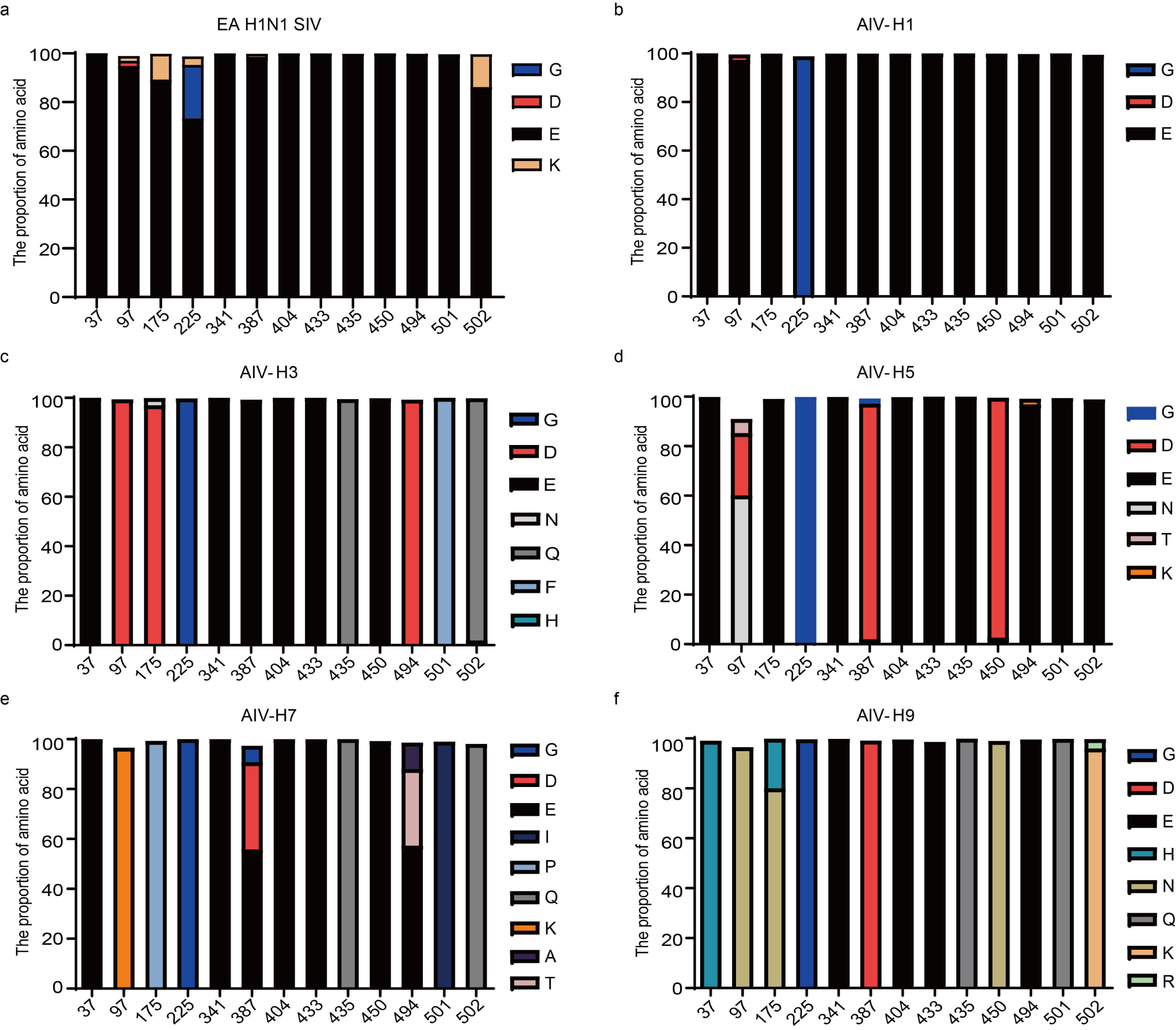
Conservation analysis of the identified carboxylation modification sites in the HA of the different IAV subtypes. Conservation analysis of carboxylation modification sites in HA protein sequences of (a) 539 EA H1N1 SIV, (b) 666 AIV-H1, (c) 2,571 AIV-H3, (d) 4,398 AIV-H5, (e) 2,129 AIV-H7, and (f) 3,507 AIV-H9 strains downloaded from the National Center for Biotechnology Information (NCBI). X-axis: the 13 distinct carboxylation modification sites that exclusively identified in HA proteins expressed in virus-infected WT cells.

## Notes

### Competing Interest Statement

The authors have declared no competing interest.

